# The first finding of lichen *Solorina saccata* at an algific talus slope in Korea

**DOI:** 10.1101/710947

**Authors:** Jung Shin Park, Kwang-Hyung Kim, Dong-Kap Kim, Chang Sun Kim, Sook-Young Park, Soon-Ok Oh

## Abstract

An algific talus slope is composed of broken rocks with vents connected to an ice cave, releasing cool air in summer and relatively warmer air in winter to maintain a more stable microclimate all year round. Such geological features create a very unusual and delicate ecosystem. Although there are around 25 algific talus slopes in Korea, lichen ecology of these areas had not been investigated to date. In this study, we report the first exploration of lichen ecology at an algific talus slope, Jangyeol-ri, in Korea. A total of 37 specimens were collected over 2017-2018. Morphological and sequencing analysis revealed 27 species belonging to 18 genera present in the area. Of particular interest among these species was *Solorina saccata*, as it has previously not been reported in Korea and most members of genus *Solorina* are known to inhabit alpine regions of the Northern Hemisphere. We provide here a taxonomic key for *S. saccata* alongside molecular phylogenetic analyses and prediction of potential habitats in South Korea. Sequences were generated from all *S. saccata* specimens collected in this study, together with three *S. saccata* specimens from China for comparison. Phylogenetic analysis based on nuclear small subunit (nuSSU) showed that all the *S. saccata* specimens are tightly grouped into one clade with high support values (P=0.99) and showed close relatedness to *S. spongiosa* than *S. crocea*. Additional analyses were carried out using concatenated sequences: 65 sequences of mitochondrial small subunit (mtSSU) combined with nuLSU and 66 sequences of RPB1 with nuLSU. In these analyses, genus *Peltigera* was found to be the closest genus to *S. saccata*. Furthermore, regions in South Korea potentially suitable for *Solorina* spp. were predicted based on climatic features of known habitats around the globe. Our results showed that the suitable areas are mostly at high altitudes in mountainous areas where the annual temperature range doesn’t exceed 26.6°C. Further survey of other environmental conditions determining the suitability of *Solorina* spp. should lead to a more precise prediction of suitable habitats and trace the origin of *Solorina* spp. in Korea.

## Introduction

Algific talus slopes are geological features in high altitude areas, composed of broken rocks with vents connected to an ice cave. The air flow through these vents create a meteorologically unique area, where cold air blows out of the vents or “wind-holes” in summer and relatively warmer air is released in winter [1].

During the last ice age in the Pleistocene epoch of the Cenozoic Era, vegetation in the Northern Hemisphere spread southwards. It was reported that due to the special microclimate created by the temperature-stabilizing effect of algific talus slopes, they sheltered various migrating animals and vegetation during interglacial periods [1–3]. Plants, mite-like insects or snails typically inhabiting higher latitudes were found at algific slopes [4, 5], suggesting that these geological features provide a distinctive environment from the surrounding area.

Algific talus slopes are located at latitudes of 35-45° north [6–11]. They are found in the Northern Hemisphere in Europe, East Asia and the USA. In the USA alone, more than 400 locations have been reported in Minnesota, Iowa, Wisconsin, Illinois and Pennsylvania. In Europe, several algific talus areas are distributed in mountainous regions around the Alps [12]. In central Honshu, Japan, there are more than 80 algific talus areas. Although a number of comprehensive studies on climate, topography, surface geology and vegetation of algific talus areas have been carried out [13–16], only a few studies exist on lichen ecology and distribution in these areas.

In a survey of lichen at White Pine Hollow State Park, Iowa, USA, 71 of 117 samples were collected from algific talus areas and 4 out of 13 newly discovered species were found in these areas. Species such as *Peltigera ponojensis*, *Peltigera membranacea*, and *Physconia muscigena* were found at higher altitude than in Iowa, but were found in the field [17]. In another study, a survey of lichens was conducted in the Spruce Creek Ice Caves, Huntingdon County, Pennsylvania, USA. Many of the lichens identified including *Arctoparmelia centrifuga*, *Cladonia coccifera*, *Cladonia rangiferina*, *Porpidia tuberculosa*, *Protothelenella corrosa*, *Rhizocarpon subgeminatum*, *Stereocaulon glaucescens*, *Vulpicida pinastri* are typically native of the northern alpine habitats [7].

In South Korea, 25 algific talus slopes are distributed from Jeju Island to Gangwon province (Fig. 1) [2]. Among these areas, vegetation of Jangyeol-ri algific talus slope, in Jeongseon, Kangwon province, was dominated by plant species *Astilboides tabularis* and *Pedicularis resupinata*, which are known to be highly vulnerable to climate change, suggesting that the biota typical of the northern regions are specifically present in the area [2].

**Fig 1.**
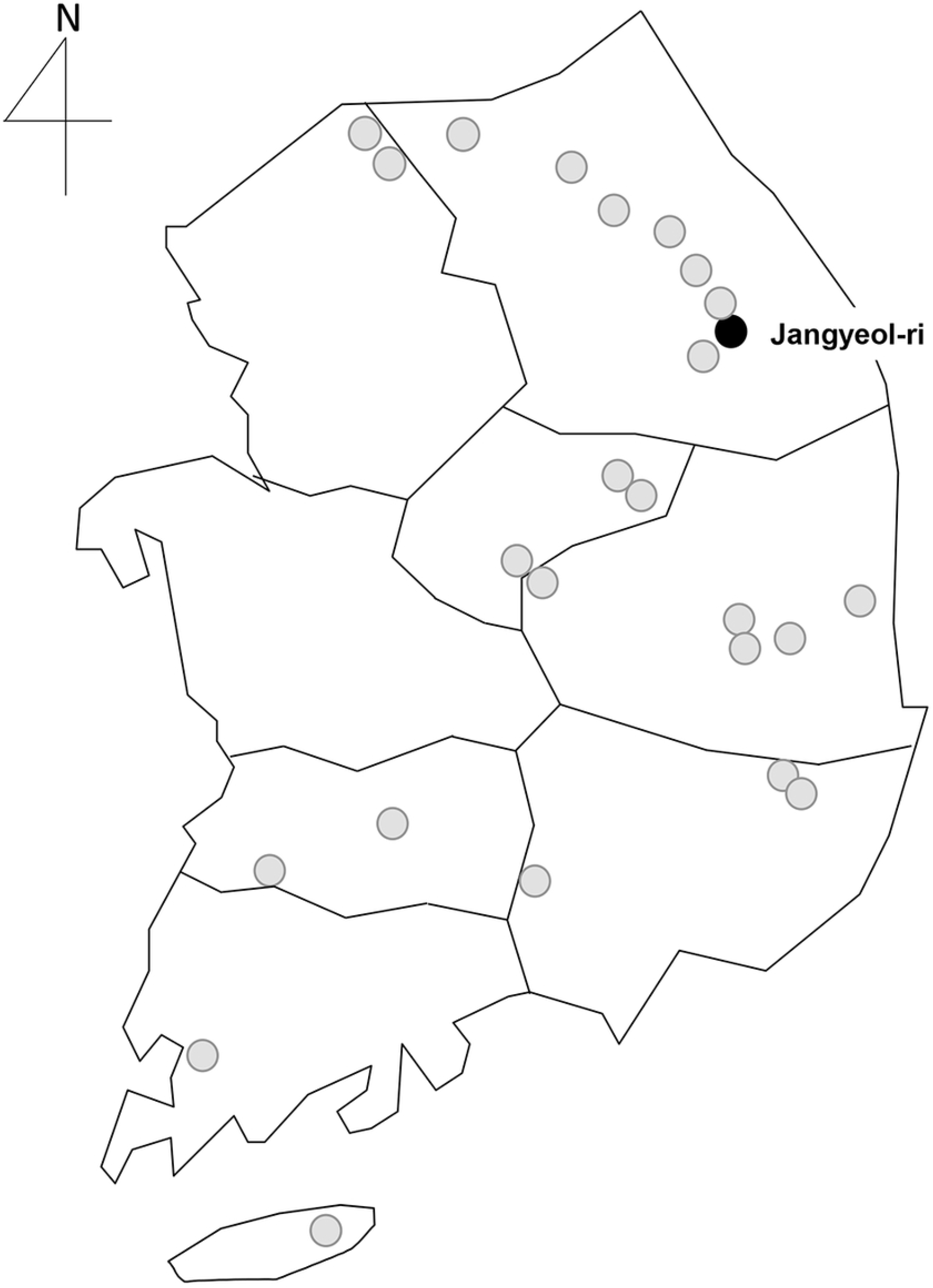
Location of 25 algific talus slopes in Korea. The black circle indicates Jangyeol-ri algific talus slope. Other sites are indicated in grey circles.

The lichen genus *Solorina* is known to be distributed in bipolar, boreal, and arctic-alpine environments [5]. The morphological characteristics of the genus *Solorina* Ach. are said to include terricolous lichens with a foliose thallus and ascoma impressed in the upper surface [18]. Genus *Solorina* belongs to the family Peltigeraceae, as is genus *Peltigera*, and is also closely related to genus *Nephroma*. The relationship between the two genera is supported by similar ascomatal ontogeny and ascus morphology [7, 8]. Unlike genus *Peltigera*, genus *Solorina* have laminal apothecia, pigmented and verrucose spores [9].

In a molecular phylogenetic analysis, genus *Solorina* was used as outgroups to assess the specificity of lichen-forming fungi and genus *Nostoc* in the genus *Peltigera* section Polydactylon [10]. The genera of Peltigeraceae including *Solorina* spp. have been used to identify the phylogenetic location of lichen symbiotic members and to improve the classification and phylogeny of Lecanoromycetes [11, 12]. In addition, genus *Solorina* has been classified as a member of family Solorinaceae, and the relationship between species is paraphyletic and monophyletic close to Peltigeraceae in a study to identify the morphogenic phylogenetic location of Peltigeraceae or Pertigerales [19]. Another study on Peltigeraceae claimed that genus *Solorina* belongs to the monophyletic family Peltigeraceae, and closely related to the sister genus *Peltigera* [9].

Unfortunately, genus *Solorina* has been largely disregarded in molecular phylogenetic studies on Peltigeraceae over the last decade have only been referred to as a sister genus to *Peltigera*. Genus *Solorina* consists of about 10 species distributed throughout the world, with 4 species in Japan and 5 species in China [14, 15], but had so far not been found in Korea.

The objectives of this study were (i) to survey lichen species in Jangyeol-ri algific talus slope, (ii) to identify the collected lichen species, and (iii) to identify lichen genus *Solorina* at morphological and molecular levels. In this study, we catalog the variety of lichen species present in Jangyeol-ri algific talus slope and also report the first finding of *Solorina saccata*, a northern lichen species, in Korea.

## Materials and Methods

### Morphological examination

All lichen specimens collected in this study were deposited at the Korean National Arboretum (KNA). Air-dried samples were observed using a stereomicroscope (Olympus SZX7) and a compound microscope (Olympus CX22LED). Water mounts were hand sectioned with a razor blade and microscopic features (ascomatal structure) were observed in water. Color reactions were conducted as described [20].

### DNA extraction and PCR amplifications

Four representative *Solorina saccata* specimens were selected and used for further molecular analyses. Lichen thalli with apothecial discs were mainly used for DNA extraction. Samples were ground and extracted with DNeasy plant mini kit (Qiagen, Valencia, CA, USA) according to the manufacturer’s instructions. PCR amplifications were conducted using Amplitaq DNA polymerase (ThermoFisher, Massachusetts, Watham, USA). The following primers were used for PCR amplifications: mtSSU1 and mtSSU3R for mtSSU [21]; NS17UCB, NS20UCB for nuSSU [22]; LIC24R [23] and LR7 [24] for nuLSU; fRPB2-7cf and fRPB2-11aR for RPB2 [25]. PCR conditions for nuSSU are as described in a previous study [26]. The following program was used for amplification of nuLSU: initial denaturation for 4 min at 94°C, followed by 30 cycles of 94°C for 40 s, 52°C for 40 s, 72°C for 50 s and then a final extension step at 72°C for 8 min. Amplified DNA was concentrated and purified using a PCR quick-spin PCR Product Purification Kit (INTRON Biotechnology, Inc. Sungnam City, Korea) for sequencing analysis.

### Sequence alignments and phylogenetic analysis

Obtained sequences were aligned with Clustal W ver. 1.83 [27] and edited using the Bioedit program. Based on the sequences, we selected commonly present sequence regions of homology, and excluded uncommonly detected sequences. We utilized Gblocks 0.91b server [28] for deleting ambiguously aligned regions in the concatenated alignment of nuSSU, nuLSU and RPB1. We prepared a three-locus concatenated dataset containing nuSSU, mtSSU+nuLSU and nuLSU+RPB1.

Phylogenetic analyses were conducted with MEGA 7.0 and MrBayes v. 3.2.6. Bayesian analysis was carried out on the data set using the Metropolis-coupled Makov chain Monte Carlo Method (MCMCMC) in MrBayes v. 3.2.6 [29, 30]. Best fit substitution models were estimated using in Akaike-information as implemented in jModelTest v 2.15. [31]. The TVM+I model in nuSSU, TIM3+I+G model in mtSSU+nuLSU, and JC+G model in nuLSU+RPB1 were selected. Each MCMCMC run was performed with four chains and 10 million generations. Trees were generated 1000 times and the first 25% was discarded. The remaining trees were determined by calculating a majority-rule consensus tree with posterior probabilities (PP).

In addition, we also performed Maximum likelihood (ML) analysis using MEGA 7.0 with 1000 ML bootstrap values (BP) and GTR+G model was applied. We selected outgroup for phylogenetic analysis according to previous research [32–34]. Phylogenetic trees were drawn using Figtree v. 1.4.3 [35] with Treeview X v. 0.5.0 and posterior probabilities above 0.80 (PP) were used near the bold branches.

### Estimation of optimum temperature for *Solorina* spp

Although *Solorina* species are native to polar and alpine regions, our specimens were collected at a relatively low altitude of 400-450 m above sea level. To determine climatic requirements for *Solorina* spp., we first identified all from collection sites of *Solorina* spp. from previously published literature and relevant reports, and then collected 30 years’ climate data on minimum and maximum air temperature for the identified habitats

A few criteria were applied to filter out unusable data. Since the climate in the Southern Hemisphere has different seasonal variations, data from the Southern Hemisphere were excluded. Sites with uncertain sampling location or unspecific records above district level were also excluded. Multiple sites located within 1 km radius were combined and considered as a single site, as we used a 1 km resolution grid dataset to extract corresponding climate data.

As a resut, the number of sites was narrowed down from 73 to 63 sites for subsequent climatological analysis. Coordinates (latitude and longitude) of the chosen sites were first estimated based on available site information and using the Google Maps (https://www.google.com/maps). Then monthly minimum and maximum temperature information at the identified coordinates were extracted from the WorldClim, a monthly climatology dataset provided at 1 km resolution (https://www.worldclim.org/). Average monthly minimum and maximum temperatures were calculated and plotted together with all the temperature ranges of the 63 sites (Fig. 7).

Site suitability for *Solorina* spp. in South Korea was determined based on the temperature data of 63 sites. The temperature range for each month was defined by the minimum and maximum values among the 63 sites. The temperature range values were used to identify suitable areas with similar monthly temperature variations in South Korea. If the local temperature of an area satisfies the above defined temperature range profile, it was deemed “Suitable”. Other areas were considered “Unsuitable”. Local temperature profiles to 1 km resolution of South Korea were obtained from a historical climate dataset produced by the Korean Meteorological Administration (KMA). High resolution KMA temperature datasets were produced from 75 KMA Automated Synoptic Observing System (ASOS) stations and 462 KMA Automated Weather Stations (AWS) using PRIsm-based Downscaling Estimation (PRIDE) model [36], and included daily weather data from 2001–2010.

## Results and Discussion

### Survey of lichen species in Jangyeol-ri algific talus slope

Jangyeol-ri algific talus slope is located at 200-450 m above sea level in Jeongseon, Kangwon province (37°27’06.14"N, 128°41’04.18"E). It is 200 m in breadth and sloped at 40 degrees (Fig. 2A-E). The geological features were consistent with the Paleozoic Ordovician strata, most of which were limestone and dark red forest soils [2].

**Fig. 2.**
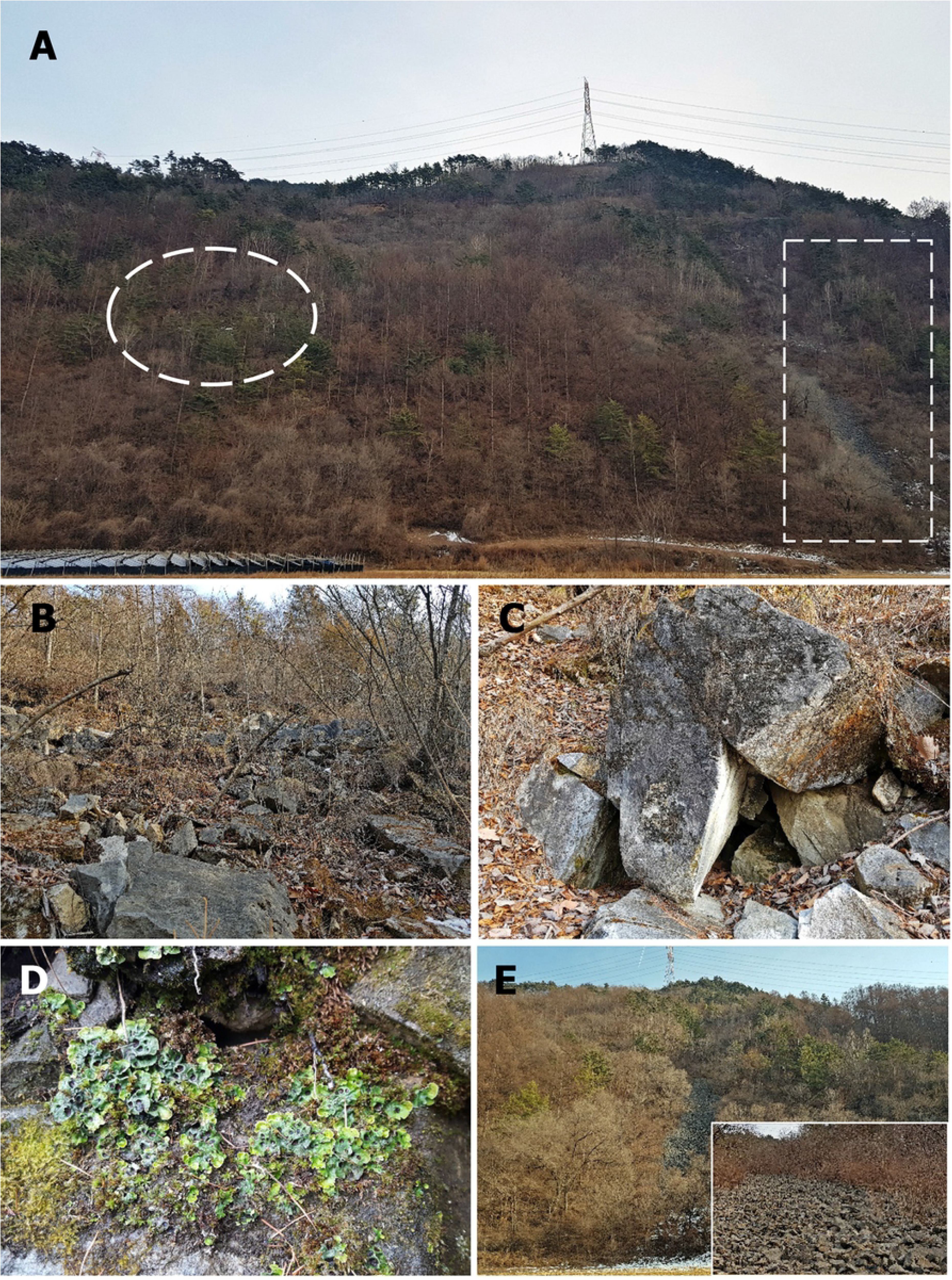
Photographs of Jangyeol-ri algific talus slope. (A) The collection area is marked by the dashed-line circle and the exposed area of the talus slope is marked by dashed-line rectangle. (B) The collection site of lichens in Jangyeol-ri algific talus slope. (C) Two vents. (D) Lichen species *Solorina saccata* in front of the vent. (E) Exposed area. The smaller insert shows more detail at the ground level.

A total of 37 lichen specimens were collected from 2017 to 2018. From these samples, 27 species belonging to 18 genera were identified by morphological examination and sequencing analysis: genus *Amandinea* (*A. punctate*); *Caloplaca* (*C. flavovirescens*); *Candelaria* (*C. concolor*); *Cladonia* (*C. furcata* subsp. *furcata* and *C. pyxidata*); *Collema* (*C. japonicum* and *C. leptaleum* var. *biliosum*); *Everniastrum* sp.; *Flavoparmelia* (*F. caperata*); *Graphis* (*G. scripta*); *Heterodermia* (*H. diademata* and *H. hypoleuca*); *Lecanora* (*L. argentata* and *L. strobilina*); *Leptogium* (*L. cyanescens*); *Myelochroa* (*M. aurulenta*); *Ochrolechia* (*O. akagiensis*); *Peltigera* (*P. elisabethae*, *P. horizontalis*, and *P. rufescens*); *Phaeophyscia* (*P. primaria*); *Protoblastenia* (*P. rupestris*); *Ramalina sp.*; *Solorina* (*S. saccata*) (Table 1).

**Table 1.**
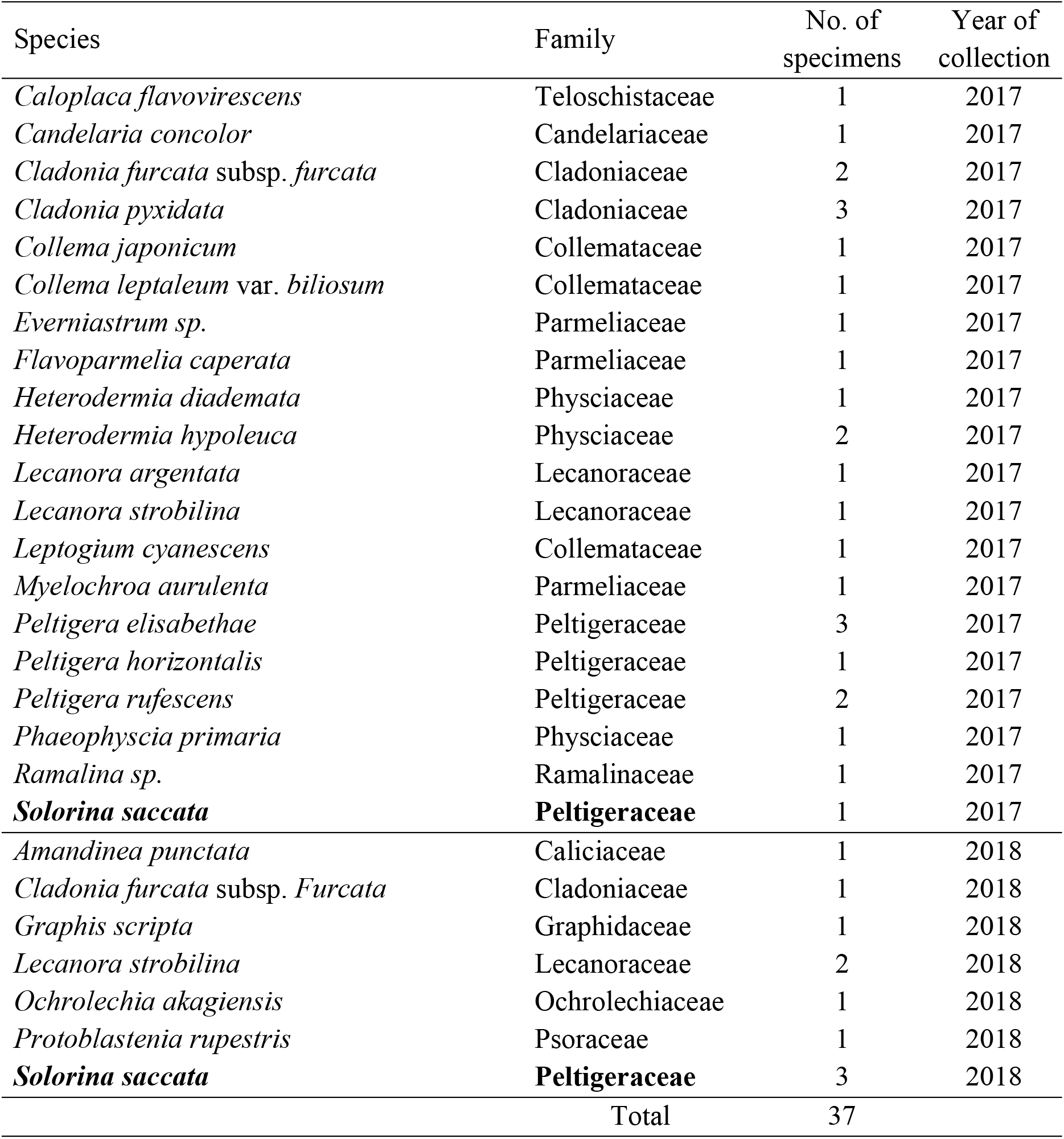
Summary of lichen species in Jangyeol-ri algific talus slope

In this survey, we could not find any lichen species overlapping with those found in previous studies on lichen inhabiting algific talus slopes in White Pine Hollow, Iowa, USA [17] and Spruce Creek Ice Cave, Pennsylvania, USA [7]. Interestingly, the most predominant lichen species we found in Jangyeol-ri algific talus belonged to genus *Solorina* (10.8% of all specimens collected), which are mainly observed in circumpolar, arctic alpine and boreal areas including Europe, Asia and North America [37]. Although there is no overlap between lichen species found in Jangyeol-ri and Spruce Creek Ice Cave, the species collected at these two different algific talus slopes were species common to northern alpine areas, indicating that the vegetation at algific talus slopes are very different from the surrounding areas at an equivalent latitude.

### Phylogenetic analysis of *Solorina saccata* using nuSSU, nuLSU and RPB1

We generated nuSSU, mtSSU, nuLSU, ITS, RPB1 and RPB2 sequences from the four *S. saccata* specimens in this study (Supplemental Table 1). Additionally, we also generated sequences from three S. saccata specimens previously collected in China for comparison. We successfully obtained sequences from all above specimens, except one Korean *S. saccata* specimen, Oh KL17-0241. The five ribosomal gene and ITS region sequences obtained from our specimens were over 99% identical to the *S. saccata* sequences in GenBank (data not shown).

For phylogenetic analysis, we retrieved reference sequences of family Peltigerales (*Dendriscocaulon*, *Nephroma*, *Leptochidium*, *Lobaria*, *Lobariella*, *Peltigera*, *Pseudocyphellaria, Solorina* and *Sticta*) from GenBank (Supplemental Table 1). A phylogenetic tree was generated using 26 nuSSU sequences (Fig. 3). All the sequences from our putative *S. saccata* specimens used in this analysis were tightly grouped into one clade with high support values (P=0.99). In addition, phylogenetic analysis revealed that *S. saccata* is more closely related to *S. spongiosa* than *S. crocea*. This is in correlation with the color of the lower surface, which is white for *S. saccta* and *S. spongiosa* and orange for *S. crocea* [38].

**Fig. 3.**
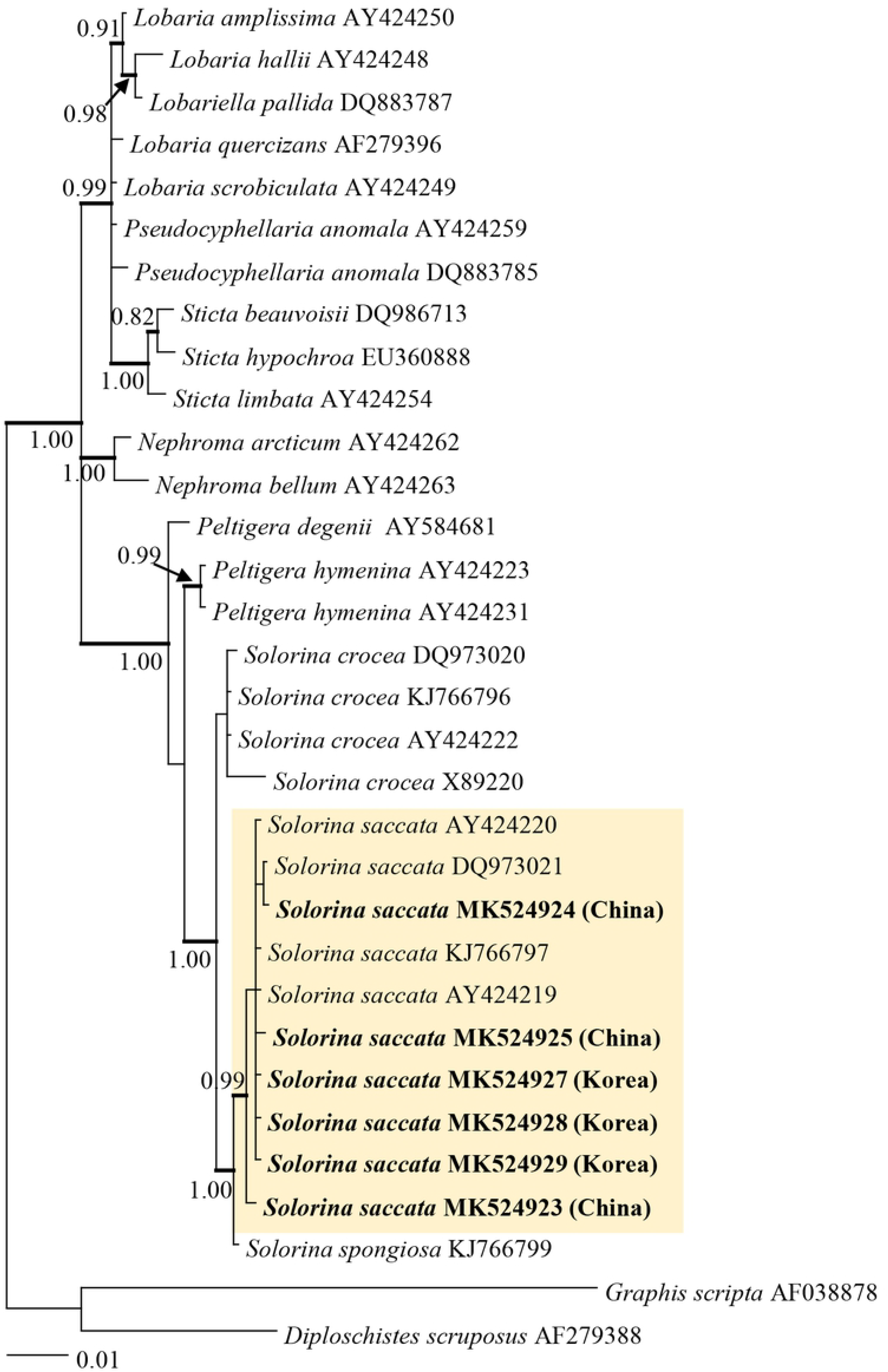
Phylogenetic relationship within family Pertigerales inferred from Bayesian analysis using nuclear small subunit (nuSSU) sequences. Only posterior probability values higher than 80% are shown. Strongly supported nodes are indicated in bold.

Furthermore, where nuSSU sequences and either mtSSU or RPB1 sequences were available, we performed phylogenetic analysis using 65 concatenated mtSSU+nuLSU sequences (Fig. 4) and 66 nuLSU+RPB1 sequences (Fig. 5). The two phylogenetic trees in Fig. 4 and Fig. 5, more convincingly showed that *S. saccata* was clearly different from other genera in the family. Within the family, genus *Peltigera* was closest to *Solorina* in both the trees from concatenated sequences. These results are in accordance with a previous analysis based on nuSSU sequences showing that *Solorina* is located in the *Hydrothyria-Peltigera* clade and is a sister-clade to *Peltigera* with 100% bootstrap support [23]. Also, when using mtSSU and nuLSU sequences, *S. crocea* and *Massalongia calrosa* were closely related to *Peltigera* with very high support values [33].

**Fig. 4.**
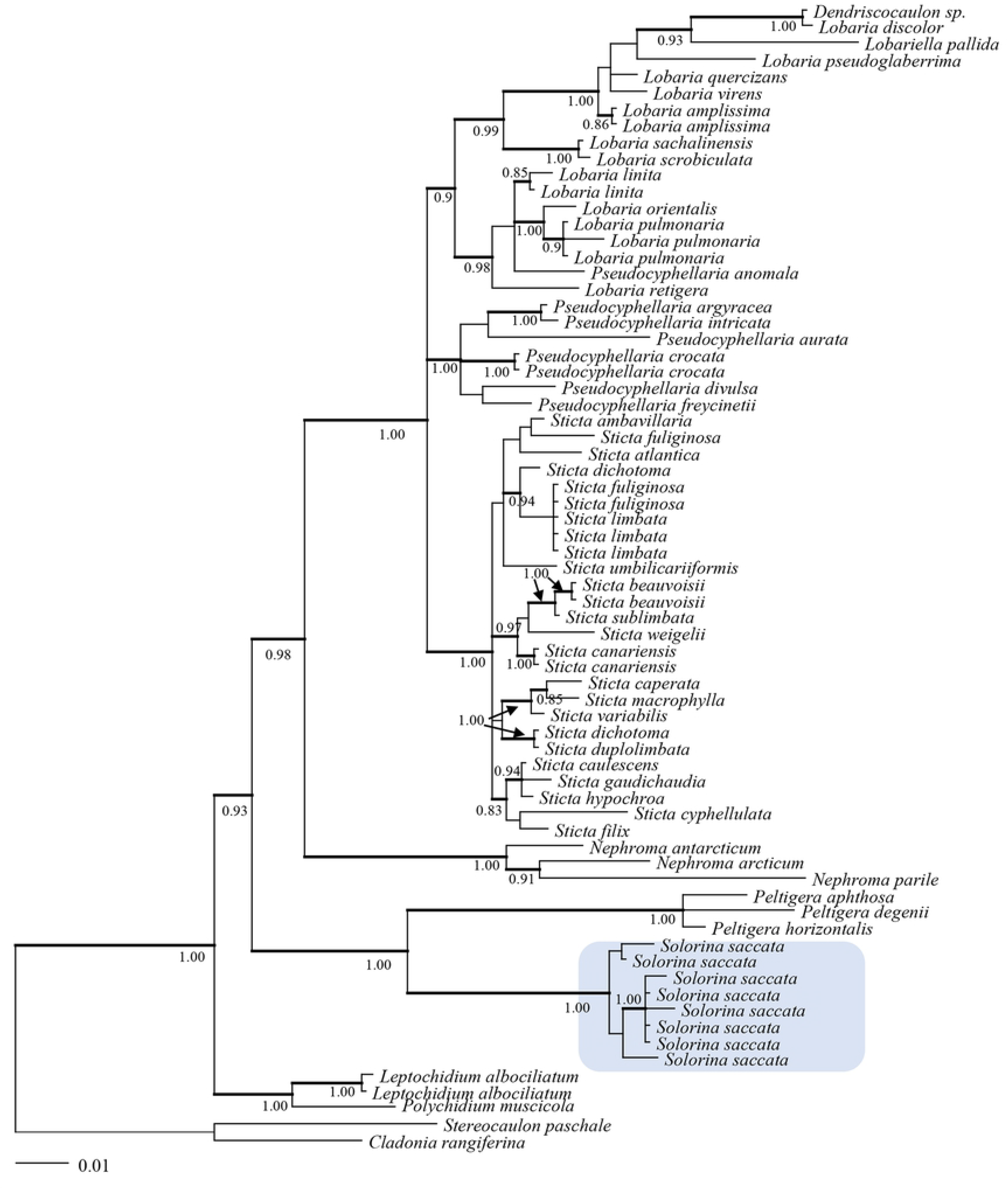
Phylogenetic relationship within family Pertigerales inferred from Bayesian analysis of concatenated mitochondrial small subunit (mtSSU) and nuclear large subunit (nuLSU) sequences. Only posterior probability values higher than 80% are shown. Strongly supported nodes are indicated in bold.

**Fig. 5.**
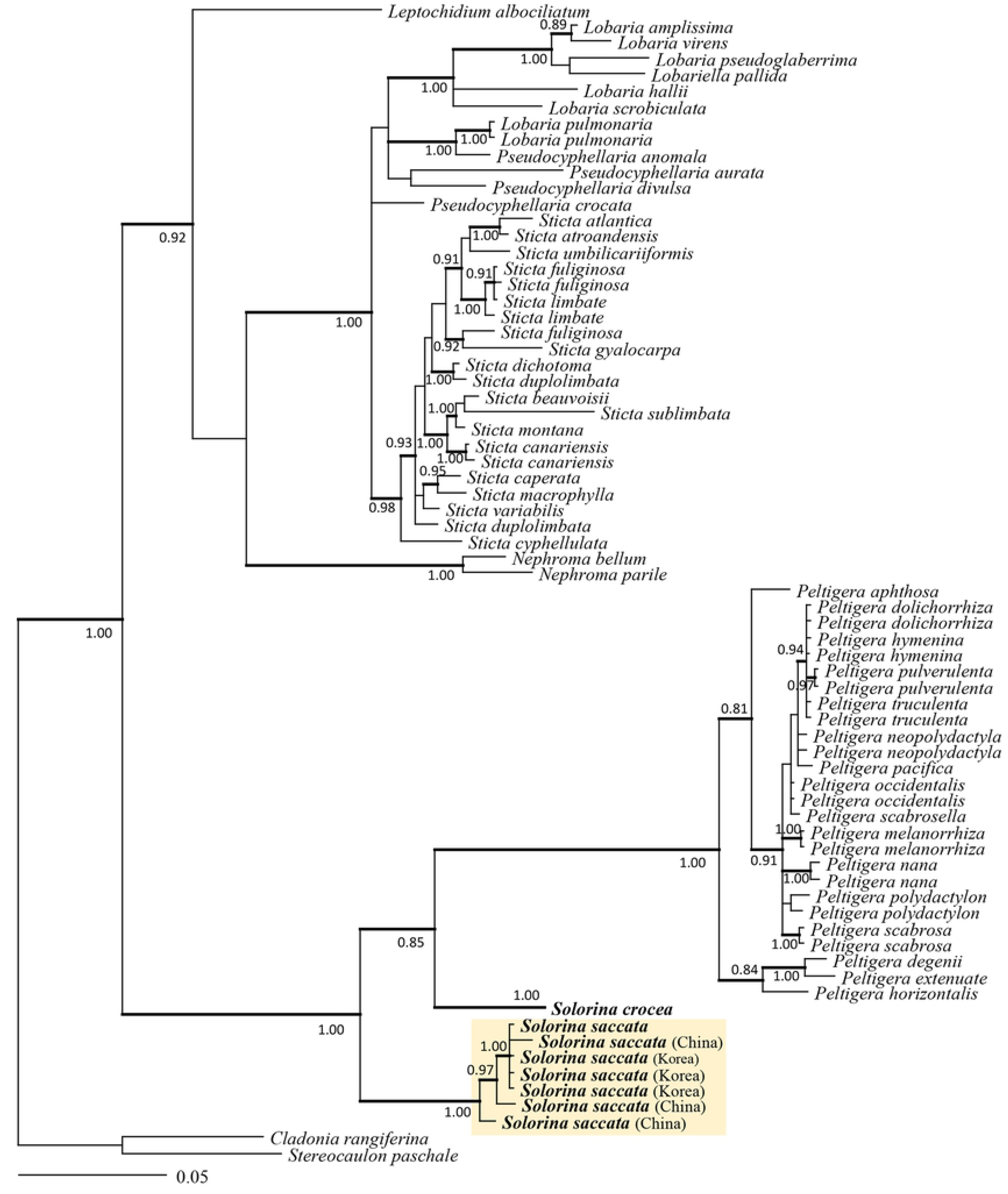
Phylogenetic relationship of Pertigerales inferred from Bayesian analysis of concatenated nuLSU and RPB1 sequences. Only posterior probability values higher than 80% are shown. Strongly supported nodes are indicated in bold.

### Comparision of average temperatures at Jangyeol-ri algific talus slope and the surrounding area, Jeongseon-gun

To investigate whether Jangyeol-ri algific talus indeed has a distinct temperature profile from the surrounding Jeongseon-gun region, we examined the monthly temperature variation (Fig. 6). Because the temperature measurements in Jangyeol-ri algific talus slope were only available from April to November 2017, the data were compared with those of Jeongseon-gun in 2017 (Fig. 6).

**Fig. 6.**
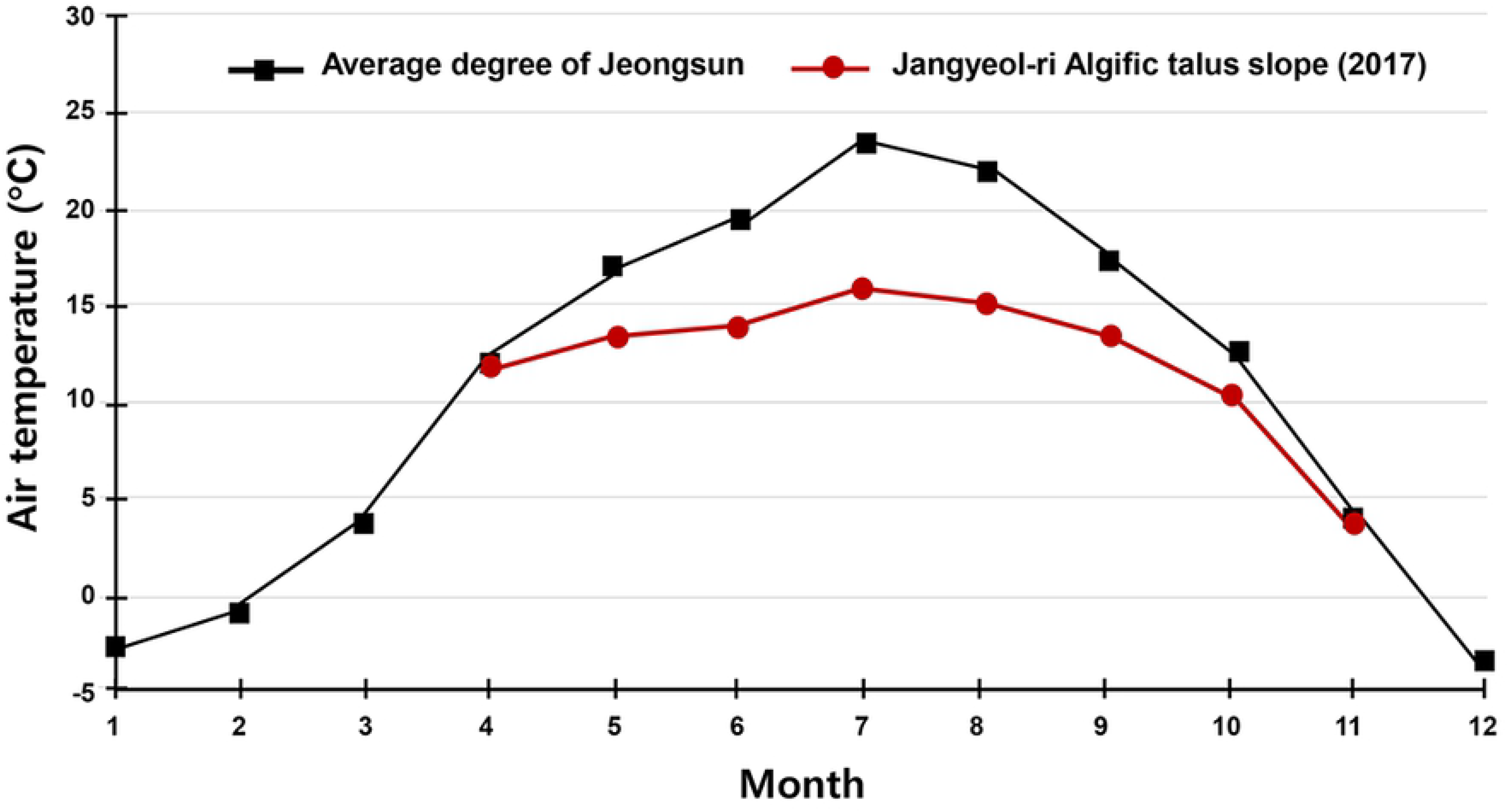
Comparison of average temperatures at the Jangyeol-ri algific talus slope and the surrounding area, Jeongsun, in 2017. X-axis: Temperature / °C, y-axis: Month.

The maximal difference in average temperature was observed in August 2017, when jangyeol-ri was 15.7°C, which was 7.3°C lower than the average temperature of 23.0°C in Jeongseon-gun. In comparison, in November 2017, the average temperature of Jangyeol-ri algific talus slope was 2.93°C, which was 0.87°C lower than the average temperature (3.8°C) in Jeongseon-gun. These results indicate that the greatest differences in temperature between Jangyeol-ri algific talus slope and the rest of Jeongsun-gun is in the summer months. It is presumed that the cooler temperatures in summer may contribute to the continued existence of *Solorina* spp. in Jangyeol-ri.

### Prediction of suitable areas for *Solorina* spp. in South Korea

Suitable areas for *Solorina* spp. in South Korea were predicted using the monthly temperature profiles of 63 sites across the globe where the *Solorina* spp. were found, assuming that monthly temperature variations are a major factor determining suitability as a habitat. Our results showed that the highest monthly maximum recorded was 26.6°C in August and the lowest monthly minimum temperature was −43.9 °C in January (Fig. 7). The minimum temperature of −43.9°C indicates that *Solorina* spp. may be able to endure the extreme cold temperatures in a state of dormancy like other lichens in polar regions.

**Fig. 7.**
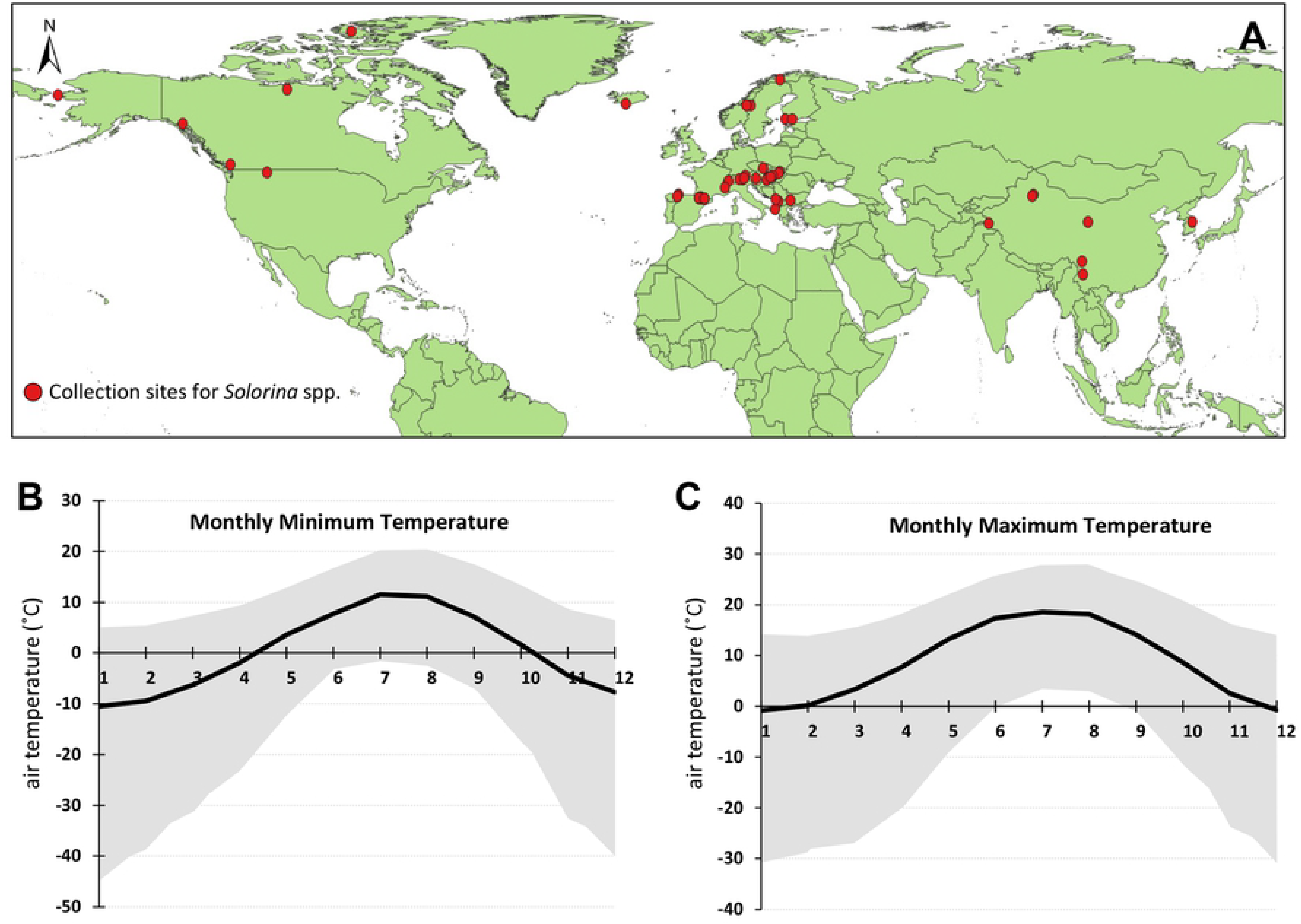
Analysis of minimum and maximum air temperatures at *Solorina* spp. habitats worldwide. (A) 63 sites where *Solorina* spp. has been reported, indicated by red dots. Monthly minimum (B) and maximum (C) temperatures of the 63 sites were extracted from the WorldClim dataset (https://www.worldclim.org/). Average (mean) monthly minimum (B) and maximum (C) temperatures are indicated as solid black lines in each graph and temperature range at the 63 sites are indicated in grey.

On the contrary, warmer temperatures may not be suitable to sustain *Solorina* spp., as previously reported areas are confined in bipolar, boreal, and arctic-alpine environments [18, 37, 39–43]. With an assumption that the *Solorina* spp. in Korea would share the same temperature requirements as those found at other sites around the world, we used the annual temperature ranges of other reported sites to predict potential areas in South Korea that *Solorina* spp. may exist.

Our result showed that the suitable areas are mostly at high altitudes in mountainous areas, where annual temperature does not exceed 26.6°C. The fact that *Solorina* spp. in Korea were found in the algific talus area, where the temperature is lower than surrounding areas in the summer months, supports our prediction results. It is highly probable that the *Solorina* spp. found in the algific talus area may not result from a migration from mountainous areas all the way down to the lower altitudes (Fig. 8). Rather, we speculate that *Solorina* spp. have been inhabiting the algific talus area likely since thousands of years ago when the algific talus area was at a high altitude before going through diastrophism.

**Fig. 8.**
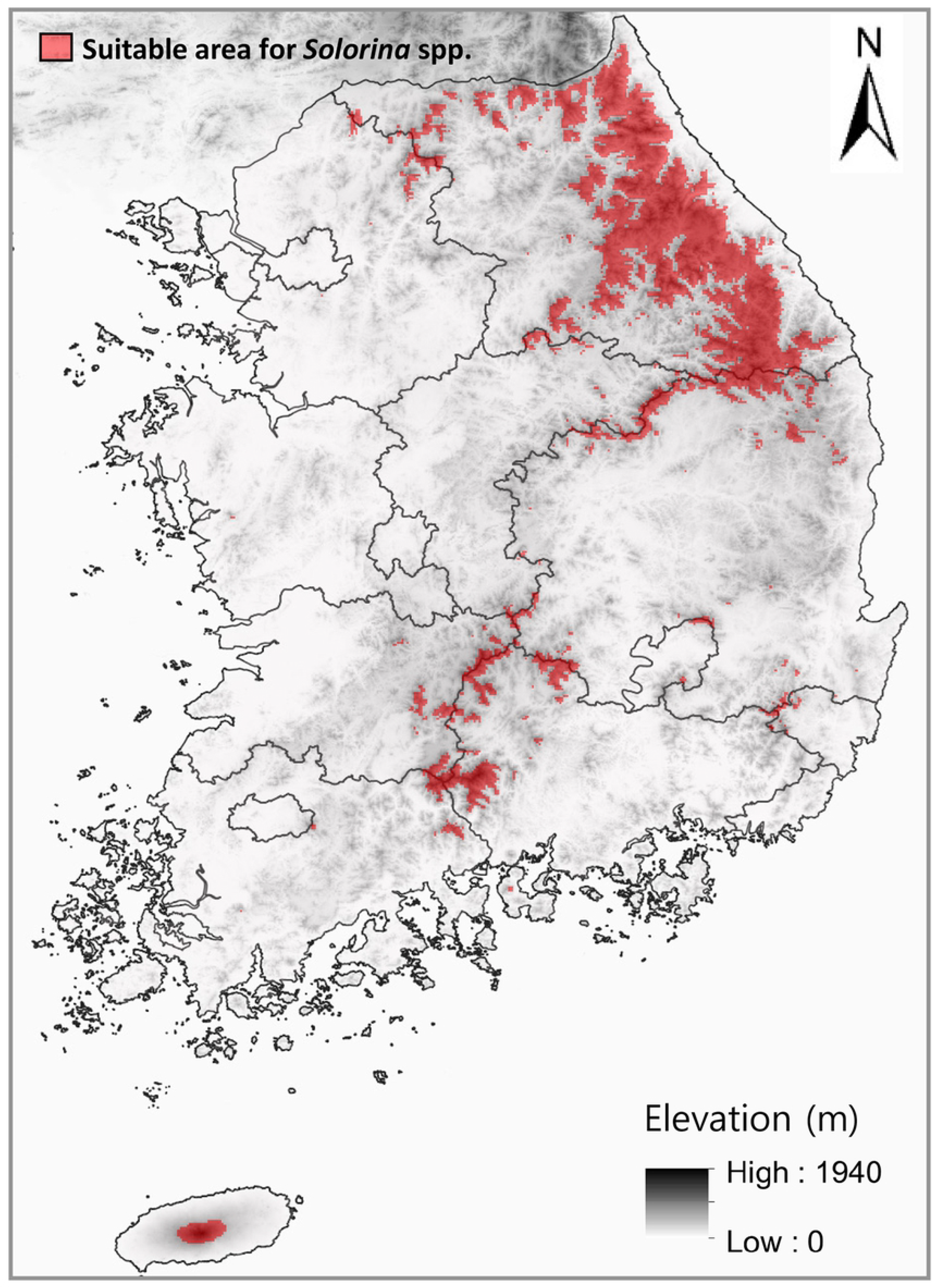
Areas potentially suitable as habitats for *Solorina* spp. in South Korea, solely based on monthly minimum and maximum temperatures in 1 km grid cells. Predicted habitats are marked in red over the elevation map of South Korea.

Other algific talus areas in Korea also show similar temperature variations to the predicted areas suitable for *Solorina* spp. Based on our preliminary analysis using temperature records of those sites over a few years, the highest temperatures recorded are generally 1-2°C less than 26.6°C, which is the highest monthly maximum temperature of the global sites of *Solorina* spp. Although it is possible that *Solorina* spp. indeed exist in the mountainous areas as predicted in our study, temperature alone may not be sufficient as an indicator of suitability for the habitat of *Solorina* spp.; other environmental factors such as relative humidity and rainfall may also be necessary determinants. However, it is very difficult to find out these environmental conditions with very limited data available from algific talus areas in Korea. Further survey on other environmental factors and their effect on *Solorina* spp. should result in more reliable prediction of their habitats in Korea and other global areas.

### Key to differentiate between the three related genera, *Peltigera*, *Nephroma* and *Solorina*

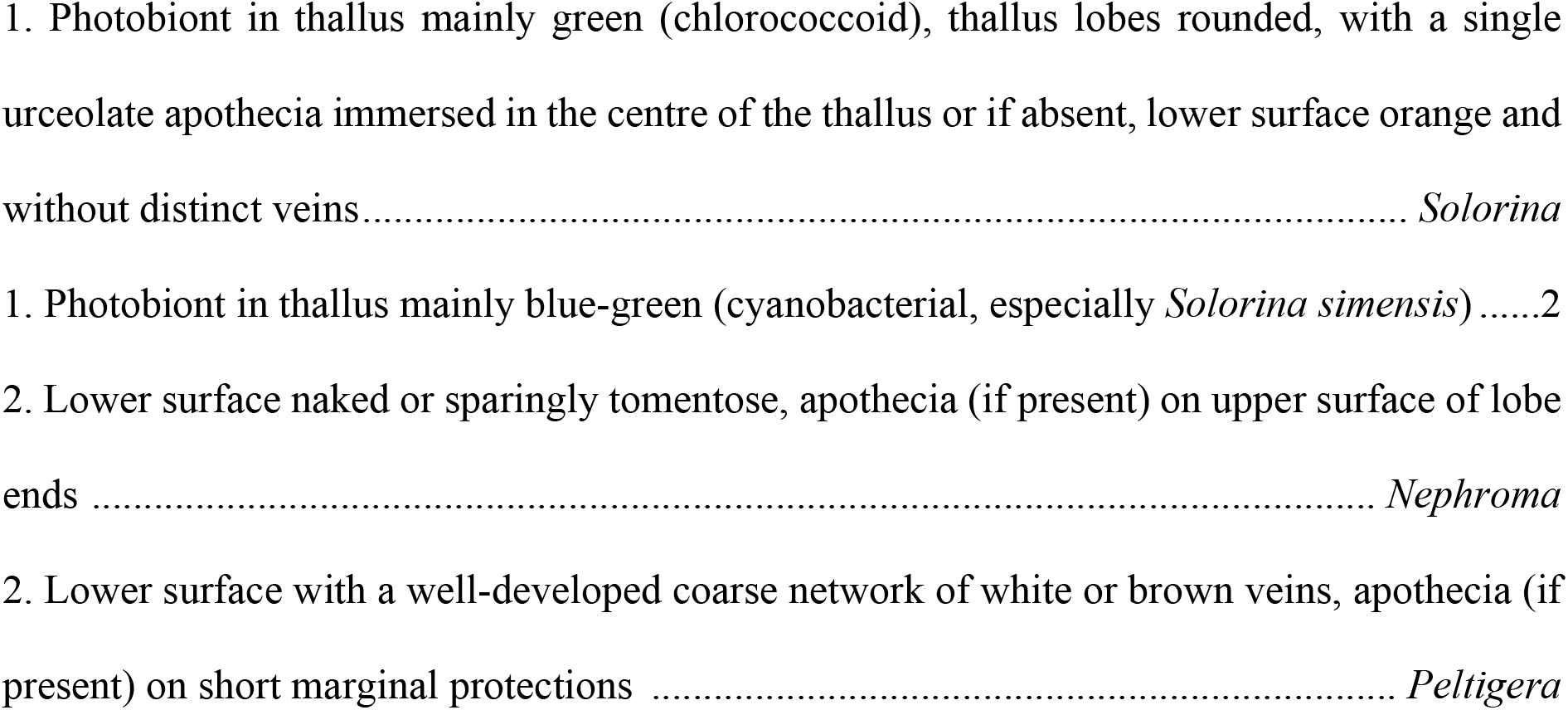

### Key to identify known species of *Solorina*

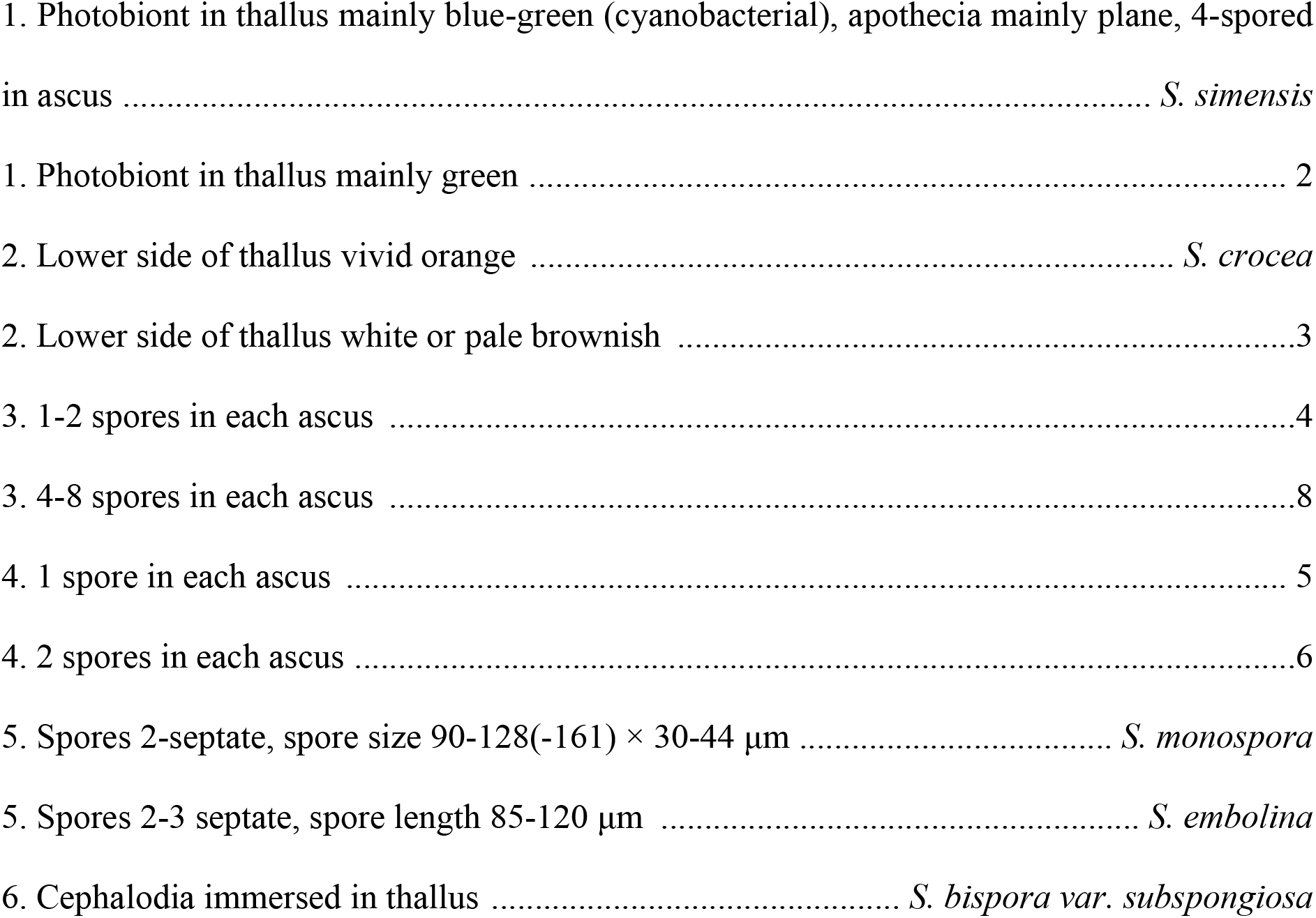
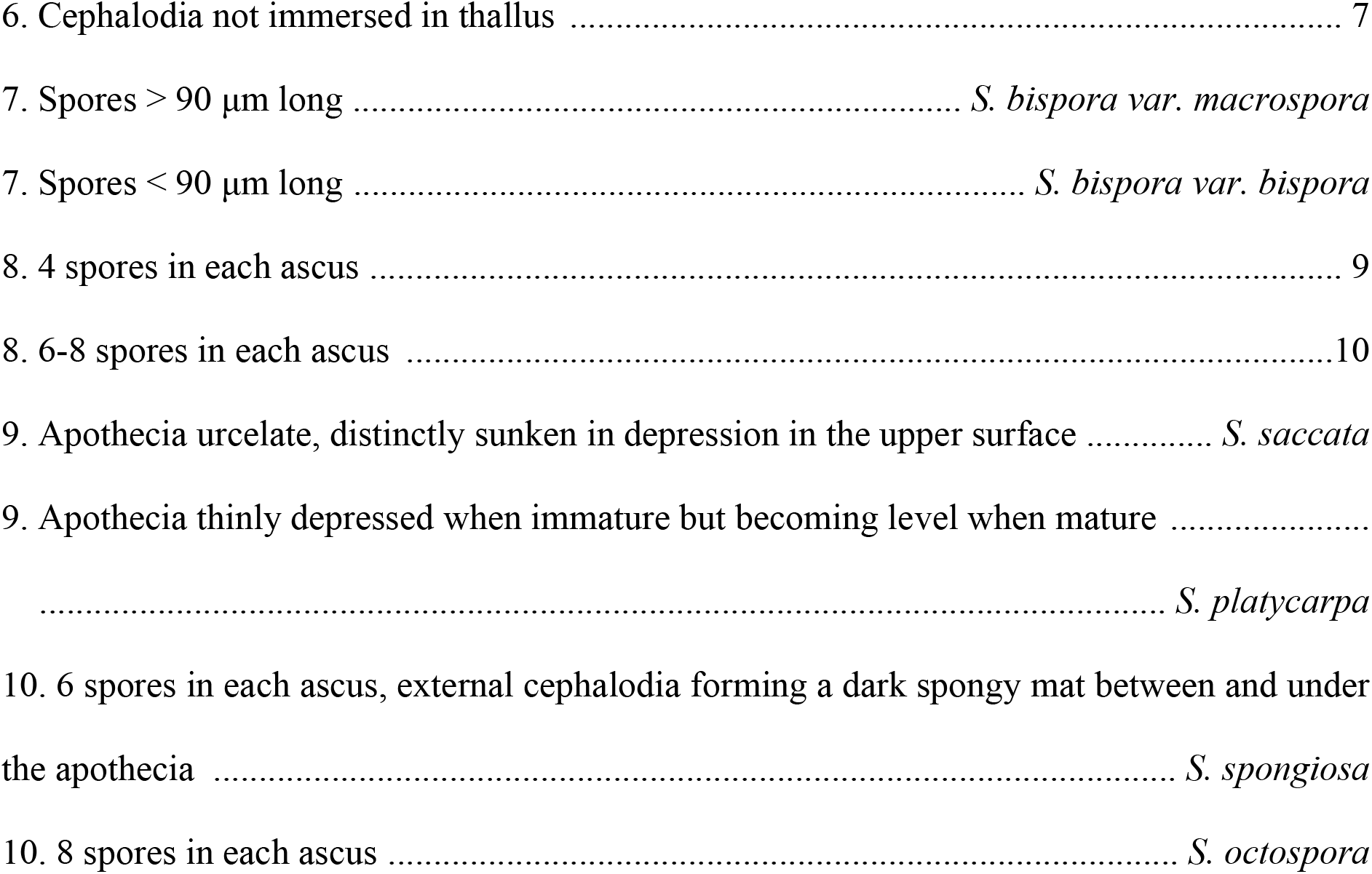

### Identification guide for *Solorina saccata*

*Solorina saccata* (L.) Ach., K. Vetensk-Acad. Nya Handl. 29: 228 (1808) (Fig. 9) Thallus well developed, spreading, forming round lobes with waxy margins, smooth, forms rosettes, greenish gray when dry but green when wet, coarse white pruinose at least close to the margin, thin spread; lower surface pale brown, rhizine sparsely present, scattered, 2-2.5 mm long, pale brown, indistinctly veined. Apothecia frequent, reddish brown, to 6.5 mm diam., deeply sunken in depression in the upper surface, with or without margin, margin 0.5 mm thick, concolor with thallus. Asci numberous, 4-spored, epihymenium brown to dark brown, 25-32.5 μm, hymenium hyaline, 155-187.5 μm; ascospores 45-53 μm × 17.5 μm, pale brown to brown, 1-septate, ellipsoid-oblong. No lichen substance detected by TLC.

**Fig. 9.**
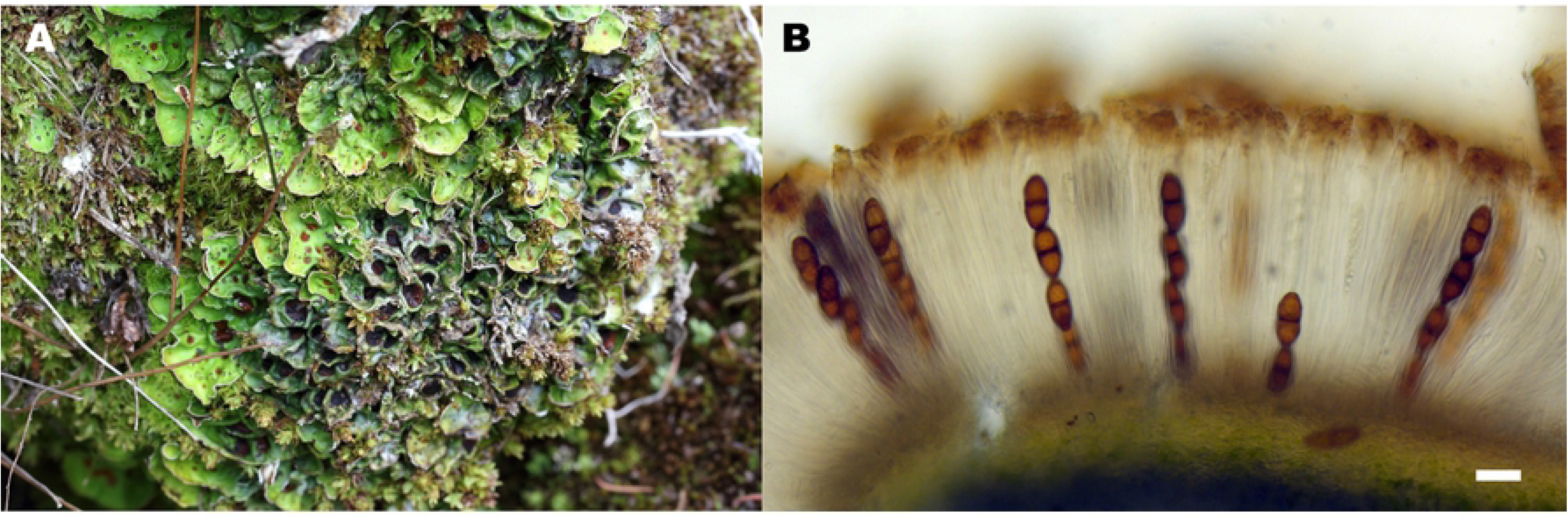
Morphology of *Solorina saccata*. (A) *Solorina saccata* KL17-241; (B) Cross section of the ascomata (4-spored asci) (Scale bars: B=40 µm).

## Acknowledgements

The authors grateful to Bun-Yeon Park for assistance in DNA sequencing, Jeong-Jae Woo and Young-Nam Kwag for helpful comments. This work was supported by a grant from the Korean Forest Service Program through the Korea National Arboretum (KNA1-1-22, 17-2), and the National Research Foundation of Korea (NRF-2017R1D1A1B04035888).

## Author Contributions

### Conceptualization

Jung Shin Park, Sook-Young Park, Soon-Ok Oh

### Data analysis

Jung Shin Park, Kwang-Hyung Kim, Dong-Kap Kim, Sook-Young Park

### Funding acquisition

Chang Sun Kim, Sook-Young Park, Soon-Ok Oh

### Investigation

Jung Shin Park, Kwang-Hyung Kim, Sook-Young Park Wring-original draft: Jung Shin Park, Kwang-Hyung Kim, Sook-Young Park, Soon-Ok Oh

## Supporting information

S1 Table 1. A total 126 taxa and the generated or retrieved nuSSU, nuLSU, mtSSU, RPB1, RPB2, and ITS sequences from NCBI.

